# Contribution of amino acids in the Active site of Dipeptidyl Peptidase 4 to the catalytic action of the enzyme

**DOI:** 10.1101/2023.07.14.549102

**Authors:** Kathrin Gnoth, Joachim Wolfgang Bär, Fred Rosche, Jens-Ulrich Rahfeld, Hans-Ulrich Demuth

**Affiliations:** Hochschule Anhalt, Department of Applied Biosciences and Process Engineering, Köthen, Germany; Boehringer Ingelheim Pharma GmbH & Co. KG, Biopharmaceuticals Cell Culture & DP, Biberach/Riß, Germany; Fraunhofer Institute for Cell Therapy and Immunology, Department of Drug Design and Target Validation, Halle, Germany

**Author notes:** Corresponding author: Kathrin Gnoth, (KG).

**Keywords:** DP4, Enzyme substrate interactions, Kinetic investigations, Mutagenesis, Proline specificity, Rate limiting step

## Abstract

Dipeptidyl peptidase 4 (DP4)/CD26 regulates the biological function of various peptide hormones by releasing dipeptides from their N-terminus. The enzyme is a prominent target for the treatment of type-2 diabetes and various DP4 inhibitors have been developed in recent years, but their efficacy and side effects are still an issue. Many available crystal structures of the enzyme give a static picture about enzyme-ligand interactions, but the influence of amino acids in the active centre on binding and single catalysis steps can only be judged by mutagenesis studies.

In order to elucidate their contribution to inhibitor binding and substrate catalysis, especially in discriminating the P_1_ amino acid of substrates, the amino acids R125, N710, E205 and E206 were investigated by mutagenesis studies.

Our studies demonstrated, that N710 is essential for the catalysis of dipeptide substrates. We found that R125 is not important for dipeptide binding but interacts in the P_1_’position of the peptide backbone. In contrast to dipeptide substrates both amino acids play an essential role in the binding and arrangement of long natural substrates, particularly if lacking proline in the P_1_ position. Thus, it can be assumed that the amino acids R125 and N710 are important in the DP4 catalysed substrate hydrolysis by interacting with the peptide backbone of substrates up- and downstream of the cleavage site.

Furthermore, we confirmed the important role of the amino acids E205 and E206. However, NP Y, displaying proline in P_1_ position, is still processed without the participation of E205 or E206.

## Introduction

Dipeptidyl peptidase IV/CD26 (DP4, EC 3.4.12.5) is an extracellular serine protease with multiple physiological functions. The protein possesses a proteolytic activity which leads to the release of dipeptides from the N-terminus of polypeptides. The enzyme hydrolyses preferentially after proline in P_1_ position but also accepts alanine or serine [1,2]. Consequently, DP4 has the potential to modulate cytokines, chemokines, neuropeptides, and growth factors containing these residues at the N-terminal penultimate position [3–7]. Furthermore, DP4 is implicated to be involved in several physiological processes by binding proteins such as adenosine deaminase, collagen, fibronectin, HIV coat protein gp120, plasminogen, HIV tat protein and the tyrosine phosphatase, CD45. However, these protein interactions do not affect the proteolytic activity of the enzyme [8–11].

DP4 is present on the surface of stimulated T cells, B cells and natural killer cells [12]. Additionally, it is found in a variety of tissues on epithelial, endothelial and acinar cells. In this context, the usage of the designation “CD26” refers to the discovery and its localisation as a cell surface antigen of lymphocytes. Within the hematopoietic system, the T cell activation marker, CD26, is an important factor in T cell-mediated immune response by modifying the activity of peptides involved in immune regulation [13].

The incretins, glucagon-like peptide 1 (GLP-1_7-36_) and glucose-dependent insulinotropic peptide (GIP) are substrates of DP4 [14]. These gastrointestinal hormones stimulate the glucose-dependent insulin secretion [15,16]; inhibit glucagon release and gastric emptying and enhance growth and differentiation of ß-cells and insulin gene expression [17–19]. Inhibition of DP4 in wild type and diabetic mice leads to increased levels of unprocessed GLP-1 and GIP in the circulation, enhanced insulin secretion, and improved glucose tolerance. Selective inhibitors of DP4 improve plasma glucose levels in human type II diabetics [20]. Thus, the inhibition of DP4 is a promising concept to control blood glucose homeostasis in the treatment of type 2 diabetes patients [21,22]. Hence, many orally-administered DP4 inhibitors have been developed by various companies such as sitagliptin, vildagliptin, saxagliptin and linagliptin [23]. Since these inhibitors are suffering from side effects and efficacy issues, there is still an interest in discovering new anti-diabetic drugs [24]. To meet this objective, detailed information about key features of amino acids in the active centre of DP4 and their contribution to substrate and inhibitor binding will push this issue forward. Among the multitude of described DP4 inhibitors, there are also substrate-like dipeptide derivatives [25]. The natural occurring tripeptides diprotin A (Ile-Pro-Ile) and diprotin B (Val-Pro-Leu) were also reported to have an inhibition effect on DP4 [26]. Rahfeld *et al.* [27] showed that these tripeptides are DP4 substrates and inhibit the enzyme competitively due to their slow turnover rate.

The catalytic mechanism of DP4 was extensively studied early by using small synthetic substrates. It was found, that the substrate hydrolysis is strict stereo specific. The scissile and P_2_-P_1_ bonds must be in *trans* configuration [28]. The P_2_, P_1_, and P_1_’ amino acid residues have to be in L-configuration. The rate limiting step for the DP4 catalysed hydrolysis of dipeptide derivates was found to be the deacylation reaction for proline substrates (proline in P_1_ position), while it is the acylation reaction for alanine or serine substrates (alanine or serine in P_1_ position) [29].

In 2003, the first crystal structure of DP4 in complex with the inhibitor valine-pyrrolidide was published by Rasmussen *et al.* [30]. Later on, the number of available DP4-crystal structures has steadily increased. Structures of DP4 co-crystallized with substrates, show important interaction sites within the active centre of the enzyme [31,32]. The catalytic triad of DP4 consists of the residues S630, D708 and H740 and is found in a large cavity at the interface of the α/β- and the propeller domain. Typical for the catalysis mechanism of serine proteases is the formation of the so called oxanion hole which is formed by the backbone NH of Y631 and the side chain OH of Y547. Surrounding the catalytic triad are amino acids responsible for the formation of a recognition pocket which is favourable for proline residues due to its size and hydrophobic assembly. This S_1_ specificity pocket is formed by the side chains of Y666, Y662, V711, V656, Y631 and W659. The environment is perfectly suited to accept proline and amino acids with small side chains. This is also indicated by the preference of proline, alanine and to a lesser extend serine or glycine at the P_1_ position [15]. Further recognition of substrates arises between their protonated N-terminus and two negatively charged site residues of E205 and E206. Replacing one of the two glutamic acids resulted almost in a loss of activity against dipeptide substrates [33]. The peptide backbone is stabilized by two amino acids namely N710 and R125 which form interactions to the peptide bond carbonyl of the substrates or inhibitors [34]. The S_1_’ site is described as flat and not well defined and does not mediate strong substrate-enzyme interactions. Aertgeerts *et al.* 2004 [31] showed by co-crystallizing the N-terminus of NP Y that the P_2_’ lysine is facing W629 and beyond that no further interactions are described. P_1_’ and P_2_’ face the solvent when bound in the tetrahedral complex. Additionally, with its freelance in these positions no favourable amino acids are described.

Despite the fact that DP4 belongs to the family of prolyl peptidases and prefers short peptide substrates showing proline in P_1_, many of the supposed physiological DP4 substrates found in *in vitro* kinetic studies possess Ala or Ser at the penultimate position [8]. Furthermore, substrate and inhibitor binding not only depend on amino acid sequence, but also on peptide length [35]. In this study, we investigate the influence of the amino acids R125, E205, E206 and N710 which are in close orientation to the catalytic triad, on substrate hydrolysis and inhibitor binding of DP4. Furthermore, the role of the depicted amino acids on the hydrolysis of substrates with varying length and different amino acids in the P_1_ position was examined. Thus, getting a new insight in the catalytic action of DP4 by defining the amino acids of growing importance during the cleavage of substrates lacking proline in P_1_ and showing the need to characterize the substrate specificity of the enzyme in regard of the substrate length.

## Results

In order to evaluate the role of distinct amino acids beyond the catalytic triad of DP4, seven enzyme variants were generated. The amino acids R125, N710, E206 and E205 in the catalytic site of DP4 were chosen in regard to specify their role in binding and hydrolysis of substrates and inhibitors. These variants were characterized by the determination of the kinetic parameters K_m_, k_cat_, k_cat_/K_m_ and IC_50_. Further, we compared the turnover rates of the physiological DP4 substrates NP Y (proline in P_1_), GIP (alanine in P_1_) and PACAP38 (serine in P_1_) by the enzyme and its variants. A focal point of this study was to investigate the role of the P_1_ amino acid residue on the hydrolysis of natural high molecular weight substrates in comparison to synthetic low molecular weight substrates and inhibitors.

The side chains of R125 and N710 interact with carbonyl groups in the backbone of substrates and inhibitors [25,31,32,34]. Both amino acids are able to participate in a hydrogen bond network and provide flexibility in the orientation of their side chains. In order to investigate the influence of these side chains, both amino acids were mutated to alanine (R125A and N710A). Further, the variant R125K was used to examine the influence of the delocalisation of the positive charge of the side chain. N710 has been replaced by aspartic acid (N710D) to introduce a negative charge and furthermore to glutamine (N710Q) in order to extend the side chain by remaining its uncharged character.

The protonated substrate N-terminus is coordinated by two glutamic acids, E205 and E206. It has been shown in earlier studies that these glutamic acids are essential for DP4 to cleave Gly-Pro-p-nitroanilide [33]. In contrast to these studies we focussed on the characterization with inhibitors and substrates of varying length (derivatives of dipeptides, tripeptides and natural peptide substrates) and different amino acids in P_1_ position. Both glutamic acids were replaced by alanine (E205A and E206A), consequently removing the negative charge of the side chain.

### Characterization with derivatives of dipeptides

Dipeptides, coupled with a C-terminal fluorophore like 7-Amido-4-Methlycoumarin (AMC) are used as standard substrates to kinetically characterize DP4. The determination of the kinetic constants K_m_, k_cat_ and k_cat_/K_m_ for the hydrolysis of the dipeptide derivatives Ala-Pro-AMC and Ala-Ala-AMC by DP4 and its variants provides information about the amino acids interacting with the P_2_- and P_1_ position of substrates.

As shown in Table 1, the kinetic parameters of R125A and R125K were comparable with the wild type enzyme, suggesting that R125 has no significant effect on binding or processing of dipeptide derivative substrates. However, N710 plays an essential role in the cleavage of these compounds.

**Table 1.** Hydrolysis of Ala-Pro-AMC and Ala-Ala-AMC by DP4 variants compared with wild type DP4. Experimental conditions: 100 mM HEPES-buffer (pH 7.6), T = 30 °C.

| DP4-variant | Substrate | $K_m$ [M] | $k_{cat}$ [$s^{-1}$] | $k_{cat}/K_m$ [$\mu M^{-1}s^{-1}$] |
| --- | --- | --- | --- | --- |
| wild type | Ala-Pro-AMC | $(1.9 \pm 0.14) \cdot 10^{-5}$ | $32 \pm 0.9$ | $1.7 \pm 0.13$ |
| | Ala-Ala-AMC | $(5.5 \pm 0.34) \cdot 10^{-4}$ | $5.7 \pm 0.18$ | $0.01 \pm 0.0003$ |
| R125A | Ala-Pro-AMC | $(3.7 \pm 0.18) \cdot 10^{-5}$ | $48 \pm 1.0$ | $1.3 \pm 0.07$ |
| | Ala-Ala-AMC | $(7.1 \pm 0.47) \cdot 10^{-4}$ | $2.3 \pm 0.08$ | $0.003 \pm 0.0002$ |
| R125K | Ala-Pro-AMC | $(3.8 \pm 0.11) \cdot 10^{-5}$ | $37 \pm 0.5$ | $0.97 \pm 0.03$ |
| | Ala-Ala-AMC | $(1.5 \pm 0.08) \cdot 10^{-3}$ | $2.6 \pm 0.09$ | $0.002 \pm 0.0001$ |
| N710A | Ala-Pro-AMC | $(1.7 \pm 0.17) \cdot 10^{-3}$ | $0.74 \pm 0.03$ | $(4.4 \pm 0.46) \cdot 10^{-4}$ |
| | Ala-Ala-AMC | $(4.1 \pm 0.08) \cdot 10^{-4}$ | $(9.4 \pm 0.06) \cdot 10^{-3}$ | $(2.3 \pm 0.05) \cdot 10^{-5}$ |
| N710Q | Ala-Pro-AMC | $(1.2 \pm 0.14) \cdot 10^{-3}$ | $0.23 \pm 0.01$ | $(1.9 \pm 0.21) \cdot 10^{-4}$ |
|  | Ala-Ala-AMC | No hydrolysis at concentrations < 10 mM |  |  |
| N710D | Ala-Pro-AMC | $(1.2 \pm 0.09) \cdot 10^{-3}$ | $19 \pm 0.6$ | $(1.6 \pm 0.14) \cdot 10^{-2}$ |
| | Ala-Ala-AMC | $(1.8 \pm 0.29) \cdot 10^{-3}$ | $0.05 \pm 0.003$ | $(2.7 \pm 0.49) \cdot 10^{-5}$ |
| E205A | Ala-Pro-AMC | $(2.4 \pm 0.09) \cdot 10^{-4}$ | $17 \pm 0.2$ | $(7.1 \pm 0.27) \cdot 10^{-2}$ |
| | Ala -Ala-AMC | $(4.7 \pm 0.84) \cdot 10^{-3}$ | $0.15 \pm 0.015$ | $(3.2 \pm 0.64) \cdot 10^{-5}$ |
| E206A | Ala-Pro-AMC | $(3.4 \pm 0.41) \cdot 10^{-3}$ | $0.8 \pm 0.05$ | $(2.3 \pm 0.32) \cdot 10^{-4}$ |
|  | Ala -Ala-AMC | No hydrolysis at concentrations < 10 mM |  |  |

Strong effects were found for the K_m_ and k_cat_ values of all three variants, while the kinetic values measured for N710D showed the lowest deviations compared with wild type DP4. The specificity of DP4 for dipeptidic Xaa-Pro-substrates is about 10-fold higher than for substrates displaying alanine in P_1_ position [36]. Therefore, two substrates, providing proline and alanine in P_1_ position were chosen to evaluate whether the mutated amino acids are involved in the determination of this specificity. While Ala-Pro-AMC was cleaved by the three variants of N710 (N710A, N710D, N710Q), it was not possible to measure kinetic data for the release of AMC from Ala-Ala-AMC by N710Q. In comparison to the wild type enzyme, the K_m_-values for the cleavage of Ala-Pro-AMC by the variants N710A and N710Q are 100-fold decreased, while the k_cat_-values dropped about factor 30.

Since the glutamic acids E205 and E206 coordinate the protonated N-terminus in the P_2_ position, effects in binding dipeptide derivatives, tripeptides and peptide hormones will be expected. The mutant E206A is nearly catalytic inactive against dipeptide derivative substrates. We were able to measure the turnover of Ala-Pro-AMC by E206A, albeit a cleavage of Ala-Ala-AMC was not detectable. The k_cat_ value for E205A was affected as well as the K_m_ values, which increase 10-fold in comparison to the wild type enzyme.

### Characterization with various inhibitors

In order to investigate the influence of the depicted amino acids of DP4 in the binding of several inhibitors, the IC_50_ values for these compounds were determined (Table 2). We used the dipeptide mimicking molecule Ile-thiazolidide [37] (S1 Fig) in comparison to several tripeptides. Tripeptides with a penultimate proline residue like diprotin A and B has been described to be substrates for DP4, but they show an apparent inhibition in a competitive substrate assay [27]. Thoma *et al*. published a crystal structure of DP4 in complex with diprotin A, showing interactions between R125 and the C-terminal carboxyl group of the tripeptide [32]. It was speculated that this interaction is only present during binding and hydrolysis of tripeptidic substrates and may stabilize the tetrahedral intermediate by the protection of the leaving group [32]. T-Butyl-Gly-Pro-Ile, with a free carboxyl terminus, was used in comparison to t-Butyl-Gly-Pro-Ile-NH_2_, to investigate the role of R125 in the binding of tripeptides while interacting with the C-terminal carboxyl group.

**Table 2.** IC50 values of the dipeptide mimicking molecule Ile-thiazolidide and several tripeptides, measured with wild type DP4 and the generated DP4 variants. Experimental conditions: 100 mM HEPES-buffer (pH 7.6); T = 30 °C; substrate concentration (Gly-Pro-AMC) in the range of Km.

| DP4-variant | inhibitor | IC<sub>50</sub> [M] |
| --- | --- | --- |
| wild type | Ile-thiazolidide | $(1.6 \pm 0.08) \cdot 10^{-7}$ |
| | Diprotin B | $(3.7 \pm 0.13) \cdot 10^{-5}$ |
| | Diprotin A | $(2.2 \pm 0.13) \cdot 10^{-6}$ |
| | t-Bu-Gly-Pro-Ile | $(3.6 \pm 0.16) \cdot 10^{-6}$ |
| | t-Bu-Gly-Pro-Ile-NH<sub>2</sub> | $(6.7 \pm 0.14) \cdot 10^{-6}$ |
| R125A | Ile-thiazolidide | $(3.1 \pm 0.41) \cdot 10^{-7}$ |
| | Diprotin B | $(2.0 \pm 0.28) \cdot 10^{-4}$ |
| | Diprotin A | $(4.0 \pm 0.53) \cdot 10^{-4}$ |
| | t-Bu-Gly-Pro-Ile | $(4.0 \pm 2.00) \cdot 10^{-4}$ |
| | t-Bu-Gly-Pro-Ile-NH<sub>2</sub> | $(8.0 \pm 1.10) \cdot 10^{-5}$ |
| R125K | Ile-thiazolidide | $(3.0 \pm 0.16) \cdot 10^{-7}$ |
| | Diprotin B | $(1.0 \pm 0.09) \cdot 10^{-4}$ |
| | Diprotin A | $(6.5 \pm 0.32) \cdot 10^{-5}$ |
| | t-Bu-Gly-Pro-Ile | $(2.0 \pm 0.21) \cdot 10^{-4}$ |
| | t-Bu-Gly-Pro-Ile-NH<sub>2</sub> | $(5.3 \pm 0.29) \cdot 10^{-5}$ |
| N710A | Ile-thiazolidide | $(4.0 \pm 1.00) \cdot 10^{-4}$ |
|  | Diprotin B | > 10 mM |
|  | Diprotin A | > 10 mM |
|  | t-Bu-Gly-Pro-Ile | > 10 mM |
| | t-Bu-Gly-Pro-Ile-NH<sub>2</sub> | $(1.4 \pm 0.53) \cdot 10^{-4}$ |
| N710Q | Ile-thiazolidide | $(2.0 \pm 0.50) \cdot 10^{-3}$ |
|  | Diprotin B | No inhibition |
| | Diprotin A | $(5.1 \pm 2.70) \cdot 10^{-3}$ |
|  | t-Bu-Gly-Pro-Ile | > 10 mM |
|  | t-Bu-Gly-Pro-Ile-NH<sub>2</sub> | > 10 mM |
| N710D | Ile-thiazolidide | $(3.0 \pm 0.15) \cdot 10^{-5}$ |
| | Diprotin B | $(2.8 \pm 3.10) \cdot 10^{-3}$ |
| | Diprotin A | $(1.1 \pm 0.40) \cdot 10^{-3}$ |
| | t-Bu-Gly-Pro-Ile | $(6.0 \pm 0.46) \cdot 10^{-4}$ |
| | t-Bu-Gly-Pro-Ile-NH<sub>2</sub> | $(4.0 \pm 0.58) \cdot 10^{-4}$ |
| E205A | Ile-thiazolidide | $(2.2 \pm 0.06) \cdot 10^{-5}$ |
| | Diprotin B | $(4.1 \pm 2.10) \cdot 10^{-3}$ |
| | Diprotin A | $(1.4 \pm 0.10) \cdot 10^{-3}$ |
| | t-Bu-Gly-Pro-Ile | $(1.7 \pm 0.10) \cdot 10^{-3}$ |
| | t-Bu-Gly-Pro-Ile-NH<sub>2</sub> | $(1.3 \pm 0.05) \cdot 10^{-3}$ |
| E206A | Ile-thiazolidide | > 10 mM |
|  | Diprotin B | > 10 mM |
|  | Diprotin A | > 10 mM |
|  | t-Bu-Gly-Pro-Ile | > 10 mM |
|  | t-Bu-Gly-Pro-Ile-NH<sub>2</sub> | > 10 mM |

The IC_50_ values of the dipeptide mimicking inhibitor Ile-thiazolidide were comparable between R125A, R125K and wild type DP4 (Table 2). These data confirmed our findings for dipeptide derivative substrates, indicating that R125 has no important influence on the binding of the amino acid residues in P_1_- and P_2_ position. However, we observed an effect for the variants R125A and R125K in the IC_50_ values of tripeptides, suggesting that the interaction between R125 and the C-terminal carboxyl group in P_1_’position is important for the binding of these compounds.

In contrast to R125, the kinetic results of the variants from N710 revealed, that this amino acid is essential for binding both, tripeptides and the dipeptide analogue Ile-thiazolidide (Table 2). There was no inhibition of the DP4 variant E206A, while we were able to measure these data for E205A. This further demonstrates that E206 is apparently more important for the coordination of the N-terminus than E205.

### Characterization with diprotin B as substrate

As described above, tripeptides with proline in P_1_ position are substrates of DP4 with low turnover rate, maybe due to the stabilisation of the tetrahedral intermediate by interaction of R125 with the C-terminal carboxyl group.

We applied a LC-MS/MS assay to quantify the hydrolysis products of tripeptides and longer peptides in a time-dependent manner and directly assess the kinetic constants without using chromophore or fluorophore compounds. To evaluate the role of R125 in tripeptide binding and hydrolysis, we measured the kinetic constants K_m_ and k_cat_ for the cleavage of diprotin B by the variant R125A in comparison to the wild type enzyme (Table 3). As shown in Table 3, the k_cat_ value of the variant is comparable with the wild type enzyme whereas the K_m_ value increased, suggesting that an interaction between R125 and the C-terminal carboxyl group of tripeptides does not decelerate the rate-limiting step.

**Table 3.** Kinetic data of the cleavage of diprotin B by DP4 and the DP4-variant R125A. Experimental conditions: T = 30 °C; 100 mM Tris-HCl, pH 7.6; substrate concentration 25 µM

| DP4-variant | $K_m$ [M] | $k_{cat}$ [s<sup>-1</sup>] | $k_{cat}/K_m$ [µM<sup>-1</sup>s<sup>-1</sup>] |
| --- | --- | --- | --- |
| wild type | $(7.2 \pm 0.58) \cdot 10^{-6}$ | $7.0 \pm 0.13$ | $0.97 \pm 0.08$ |
| R125A | $(3.7 \pm 0.26) \cdot 10^{-4}$ | $7.7 \pm 0.13$ | $(2.1 \pm 0.15) \cdot 10^{-2}$ |

### Characterization with peptide hormones

Besides characterizing the depicted amino acids with short substrates and inhibitors, their role in binding and processing of substrates with prolonged chain length was examined. The physiological substrates NP Y (proline in P_1_ position), GIP (alanine in P_1_ position) and PACAP38 (serine in P_1_ position) were chosen to investigate the role of different amino acids in P_1_ position. First, MALDI-TOF mass spectrometry analysis was used to examine whether a cleavage of these substrates was carried out by the DP4 variants (Table 4).

**Table 4.** Hydrolysis of the peptide hormones NP Y, GIP and PACAP38 by DP4-variants in comparison to the DP4 wild type enzyme. +++ hydrolysis is comparable with the data obtained from the wild type enzyme - hydrolysis below 0,01% of the wild type enzyme Experimental conditions: T = 37 °C; 100 mM Tris-HCl, pH 7.6; substrate concentration 25 µM.

| DP4-variant | Hydrolysis of NP Y<br>(H<sub>2</sub>N-Y-P-S...) | Hydrolysis of GIP<br>(H<sub>2</sub>N-Y-A-E...) | Hydrolysis of PACAP38<br>(H<sub>2</sub>N-H-S-Q...) |
| --- | --- | --- | --- |
| Wild type | +++ | +++ | +++ |
| R125A | ++ | + | - |
| R125K | +++ | ++ | - |
| N710A | + | - | - |
| N710Q | + | - | - |
| N710D | ++ | - | - |
| E206A | + | - | - |
| E205A | ++ | - | - |

The kinetic data presented in Table 4 indicate that the depicted amino acids in the active centre of DP4 have a different relevance during the processing of peptide substrates. Depending on the amino acid in P_1_ position, NP Y was still processed by the different DP4 variants. A cleavage of GIP was not observed by the variants E205A, E206A, N710A, N710Q and N710D. This demonstrates that N710, E205 and E206 are essential for the truncation of substrates with alanine in P_1_ position. The hydrolysis of GIP catalysed by the DP4 variants R125A and R125K was detectable, but at a lower rate in comparison to the cleavage of NP Y. Interestingly, none of the DP4 variants was able to process PACAP38 with serine in P_1_ position.

To investigate whether the observed effects on the substrate hydrolysis are related to differences in substrate binding or catalysis, we used the LC-MS/MS method to determine the K_m_ and k_cat_ values (Table 5).

**Table 5.**
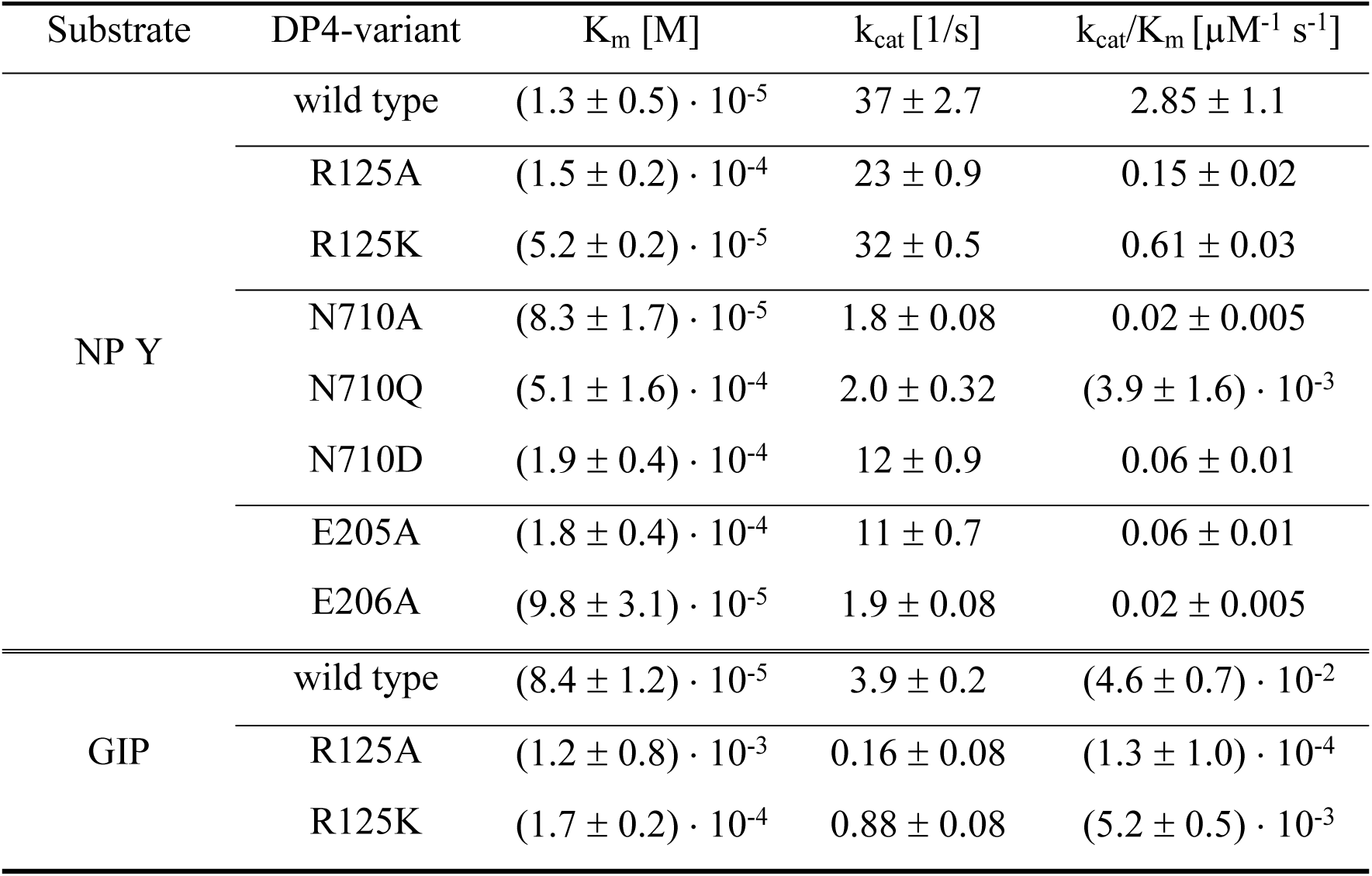
Kinetic data of the hydrolysis of NP Y and GIP by DP4 variants, compared with wild type DP4. Experimental conditions: 10 mM Tris-buffer (pH 7.6), T = 30 °C.

| Substrate | DP4-variant | K<sub>m</sub> [M] | k<sub>cat</sub> [1/s] | k<sub>cat</sub>/K<sub>m</sub> [μM<sup>-1</sup> s<sup>-1</sup>] |
| --- | --- | --- | --- | --- |
| NP Y | wild type | (1.3 ± 0.5) · 10<sup>-5</sup> | 37 ± 2.7 | 2.85 ± 1.1 |
|  | R125A | (1.5 ± 0.2) · 10<sup>-4</sup> | 23 ± 0.9 | 0.15 ± 0.02 |
|  | R125K | (5.2 ± 0.2) · 10<sup>-5</sup> | 32 ± 0.5 | 0.61 ± 0.03 |
|  | N710A | (8.3 ± 1.7) · 10<sup>-5</sup> | 1.8 ± 0.08 | 0.02 ± 0.005 |
|  | N710Q | (5.1 ± 1.6) · 10<sup>-4</sup> | 2.0 ± 0.32 | (3.9 ± 1.6) · 10<sup>-3</sup> |
|  | N710D | (1.9 ± 0.4) · 10<sup>-4</sup> | 12 ± 0.9 | 0.06 ± 0.01 |
|  | E205A | (1.8 ± 0.4) · 10<sup>-4</sup> | 11 ± 0.7 | 0.06 ± 0.01 |
|  | E206A | (9.8 ± 3.1) · 10<sup>-5</sup> | 1.9 ± 0.08 | 0.02 ± 0.005 |
| GIP | wild type | (8.4 ± 1.2) · 10<sup>-5</sup> | 3.9 ± 0.2 | (4.6 ± 0.7) · 10<sup>-2</sup> |
|  | R125A | (1.2 ± 0.8) · 10<sup>-3</sup> | 0.16 ± 0.08 | (1.3 ± 1.0) · 10<sup>-4</sup> |
|  | R125K | (1.7 ± 0.2) · 10<sup>-4</sup> | 0.88 ± 0.08 | (5.2 ± 0.5) · 10<sup>-3</sup> |

As shown in Table 5, the k_cat_ values measured for NP Y were not decreased in comparison to the wild type enzyme. In contrast to NP Y, binding as well as the rate limiting step for processing GIP was affected by the exchange of R125.

Further, a variation of the amino acids N710, E205 and E206 affected the K_m_- as well as the k_cat_ values for the hydrolysis of the physiological peptide substrates NP Y, GIP and PACAP38 (Table 5). This agrees with our findings for the dipeptide derivative substrates, indicating that these amino acids are involved in substrate binding and hydrolysis.

## Discussion

Several interactions between the active site of DP4 and its substrates/inhibitors were predicted on the basis of crystal structures of the enzyme. A crystal structure gives a static picture where the dynamics of enzyme-ligand interactions and the influence of amino acids in the active centre on catalysis and binding can hardly be judged. We performed mutagenesis studies and kinetic analysis to examine the role of amino acids around the catalytic triad in substrate catalysis and inhibitor binding.

DP4 is a member of a highly specialised protease-family, which recognizes preferentially proline in P_1_ position. It is known that amino acids such as alanine or serine can also bind in the proline specific hydrophobic S_1_ binding site. Interestingly, the change in the P_1_ position of dipeptide derivative substrates is accompanied with a change in the rate-limiting step. The deacylation reaction is the rate-limiting process for the turnover of proline substrates while it is the acylation reaction for alanine substrates. Many of the supposed physiological DP4 substrates, especially the incretins, whose prolonged half live is the main target in the treatment of human type II diabetics, possess Ala or Ser in P_1_ position. This study was performed to elucidate the contribution of amino acids in the active site beside the catalytic triad to the enzymatic specificity and the process of substrate binding and catalysis. The focus was especially on the mechanism being involved in discriminating amino acids in the P_1_ position of substrates. Further, the comparison of short synthetic substrates and longer physiological substrates should give detailed information about the relationship between substrate length and DP4 specificity.

Commonly, DP4 activity is characterized by measuring the release of a C-terminal fluorophor (e.g. AMC) or chromophor (e.g. pNA) from dipeptide derivatives. We evaluated the interactions between the catalytic side residues of DP4 and the amino acids in the P_2_- and P_1_ position by using the substrates Ala-Pro-AMC, Ala-Ala-AMC and the dipeptide mimicking inhibitor Ile-Thiazolidide respectively. Further, the use of Ala-Pro-AMC in comparison to Ala-Ala-AMC, allowed specifying the influence of the mutated side chains in the discrimination of the P_1_ amino acid in dipeptide derivatives.

To verify the influence of the investigated DP4 amino acids on the interaction with the P_1_’ position, we extended the peptide length from 2 to 3 amino acids and kinetically characterized the DP4 variants with several tripeptides. As previously described, tripeptides with a proline residue in P_1_ position are substrates of DP4 with a poor turnover rate, leading to a competitive inhibition of processing fluorogenic substrates. In order to evaluate the role of the varied amino acids, the apparent IC_50_ values for the inhibition of diprotin A, diprotin B, t-Butyl-Gly-Pro-Ile and t-Butyl-Gly-Pro-Ile-NH_2_ by the DP4 variants were determined. Further we measured the K_m_ and k_cat_ values for the cleavage of diprotin B by the variants of R125, to examine the role of R125 in tripeptide binding and hydrolysis.

The influence of the varied amino acids in binding longer physiological DP4 substrates was also examined and compared with the kinetic data obtained for dipeptide derivatives and tripeptides. The role of the amino acid in P_1_ position was evaluated by using the peptide hormones NP Y (proline in P_1_ position), GIP (alanine in P_1_ position) and PACAP38 (serine in P_1_ position). First, we examined by MALDI-TOF mass spectrometry analysis the ability of the DP4 variants to hydrolyse the peptide hormones. The k_cat_ and K_m_ values were determined by HPLC-MS/MS analytic to investigate whether the effects in substrate processing are related to substrate binding or the rate-limiting step.

### Role of N710 in substrate binding and catalysis

Crystal structures of DP4 in complex with different dipeptide mimicking inhibitors revealed, that N710 together with R125 form a polar electrophilic environment for coordinating the P_2_ carbonyl oxygen [25,30,38]. The three generated variants of N710 (N710A, N710Q and N710D) displayed strong effects in binding and hydrolysis of dipeptide derivatives and tripeptides as well as peptide hormones. The exchange of asparagine to alanine (N710A) results in a loss of the capability to form hydrogen bonds. Thus, coordination of the P_2_ carbonyl oxygen was not possible during substrate binding and hydrolysis, resulting in approximately 1000-fold decreased k_cat_/K_m_ values for dipeptide derivatives. The variant N710Q showed similar effects as N710A in substrate/inhibitor binding and processing. Exchanging asparagine to glutamine, and extending the side chain by one methyl group, keeps the ability to serve as proton donor for hydrogen bond formation. However, the interaction between glutamine in position 710 and the P_2_ carbonyl oxygen seems to be impossible probably due to steric changes. Further, we were not able to measure kinetic data for the hydrolysis of tripeptides and peptide hormones by the variants N710A and N710Q. In contrast, these substrates were cleaved by the variant N710D albeit with decreased k_cat_/K_m_ values. It can be assumed that the carboxyl group of aspartic acid in position 710 serves as proton donor for hydrogen bond formation to the substrate carbonyl oxygen instead of N710. The lengths of the side chains are comparable between asparagine and aspartic acid albeit having different proton donor properties, resulting in varied K_m_ and k_cat_ values.

The variants of N710 showed effects on K_m_, k_cat_ and IC_50_ values for substrates of all different lengths. This indicates that coordination of the P_2_ carbonyl oxygen by N710 is in general important for substrate/inhibitor binding and hydrolysis. Our investigations with substrates offering different amino acids in P_1_ position indicate the increasing importance of N710 during cleavage of substrates which possess amino acids with shorter side chains than proline (alanine or serine) in P_1_ position. In this case, further interactions, e.g. with N710, are required for substrate binding and coordination of the scissile bond resulting in its cleavage. The proline in P_1_ position of NP Y fits perfectly into the hydrophobic S_1_ pocket in the active site of DP4, whereby the participation of N710 is not essential.

### Role of R125 in substrate binding and catalysis

Co-crystallisation studies of DP4 with several dipeptidic inhibitors show, that R125 interacts with the carbonyl oxygen in P_2_ position [25,30,38] and was therefore described as a secondary substrate binding hotspot in molecular docking analysis [24]. Thoma and co-workers published a crystal structure of DP4 in complex with diprotin A, whereby describing an interaction of R125 with the C-terminal carboxyl group of the tripeptide [32]. This interaction is only present in tripeptidic substrates due to the free carboxyl-terminus and may stabilize the tetrahedral intermediate by the protection of the leaving group. Further, the crystal structure of DP4 in complex with truncated NP Y (tNP Y, residues 1-10 of NP Y) shows an interaction between R125 and the carbonyl oxygen in P_1_’position [31].

We measured similar kinetic data for the wild type enzyme and the variants of R125 in binding and hydrolysis of dipeptide derivatives and the dipeptide analogue inhibitor Ile-Thiazolidide. Thus, we concluded that the interaction between R125 and the P_2_ carbonyl oxygen described on the basis of several crystal structures has no important role in substrate/inhibitor binding or processing.

In contrast to the findings for dipeptide derivative substrates, our data show that the variants of R125 have an impaired binding ability for tripeptides. The IC_50_ values for tripeptide binding by the variants R125A and R125K were found to be increased in comparison to the wild type enzyme (Table 2). This indicates that R125 interacts with the amino acid in P_1_’position. Tripeptides possess a free C-terminal carboxyl group, which belongs to an amide bond in longer substrates. To evaluate the influence of this P_1_’ carboxyl group on tripeptide binding, we measured the IC_50_ values for t-Butyl-Gly-Pro-Ile in comparison to t-Butyl-Gly-Pro-Ile-NH_2_. The IC_50_ values dropped equally for the enzyme variants of N710, E205 and E206 when measured with both tripeptides. In contrast, the values for the binding of a tripeptide with a C-terminal carboxyl group (t-Butyl-Gly-Pro-Ile) by the variants of R125 were 100-fold decreased in comparison to the wild type enzyme (Table 2), whereas the IC_50_ values for t-Butyl-Gly-Pro-Ile-NH_2_ dropped 10-fold. In case of t-Butyl-Gly-Pro-Ile-NH_2_ a hydrogen bond between R125 and the P_1_’ carbonyl oxygen is most likely, whereas a salt bridge to the C-terminal carboxyl group could be assumed when binding t-Butyl-Gly-Pro-Ile. A salt bridge is an interaction with a higher binding energy in comparison to a hydrogen bond, explaining the differences in the impairment of the binding of t-Butyl-Gly-Pro-Ile and t-Butyl-Gly-Pro-Ile-NH_2_ by the variants of R125.

It has been hypothesised that the interaction between R125 and the C-terminal carboxyl group of diprotin A stabilizes the tetrahedral intermediate during catalysis [32]. To evaluate the effect of R125 in tripeptide binding and hydrolysis, the K_m_ and k_cat_ values were determined for the truncation of diprotin B by the variant R125A. The kinetic data show that R125 plays an important role in tripeptide binding. This was demonstrated with the increase in the IC_50_ values of all used tripeptides (Table 2) as well as the K_m_ values of diprotin B (Table 3). Interestingly, the k_cat_ value for the hydrolysis of diprotin B by R125 was comparable to the wild type enzyme. This suggests that R125 does not decelerate the rate-limiting step in the hydrolysis of tripeptides showing a proline in P_1_ position.

The cleavage of the physiological substrates NP Y, GIP and PACAP38 is also affected by the variation of R125. Interestingly, the kinetic investigations showed a strong influence of the amino acid in P_1_ position. While PACAP38 (serine in P_1_ position) was not hydrolysed, GIP (alanine in P_1_ position) was cleaved by R125A and R125K, but with a significant lower hydrolysis rate as by the wild type enzyme. Figure 1 shows, that the hydrolysis rate of GIP also depends on the interaction with the amino acid placed on position 125. K125, as well as R125, is able to form a hydrogen bond to the carbonyl oxygen in P_1_’position. The guanidino-group of arginine provides two centres to form hydrogen bonds whereas the ε-amino-group of lysine possesses only one centre. Consequently, the reduced hydrolysis velocity of GIP by R125K is probably caused by the weaker interaction between K125 and the carbonyl oxygen in P_1_’position. In contrast, alanine in position 125 has not the ability to form a hydrogen bond to the P_1_’ carbonyl oxygen, which is reflected by the very slow hydrolysis rate.

**Figure.**
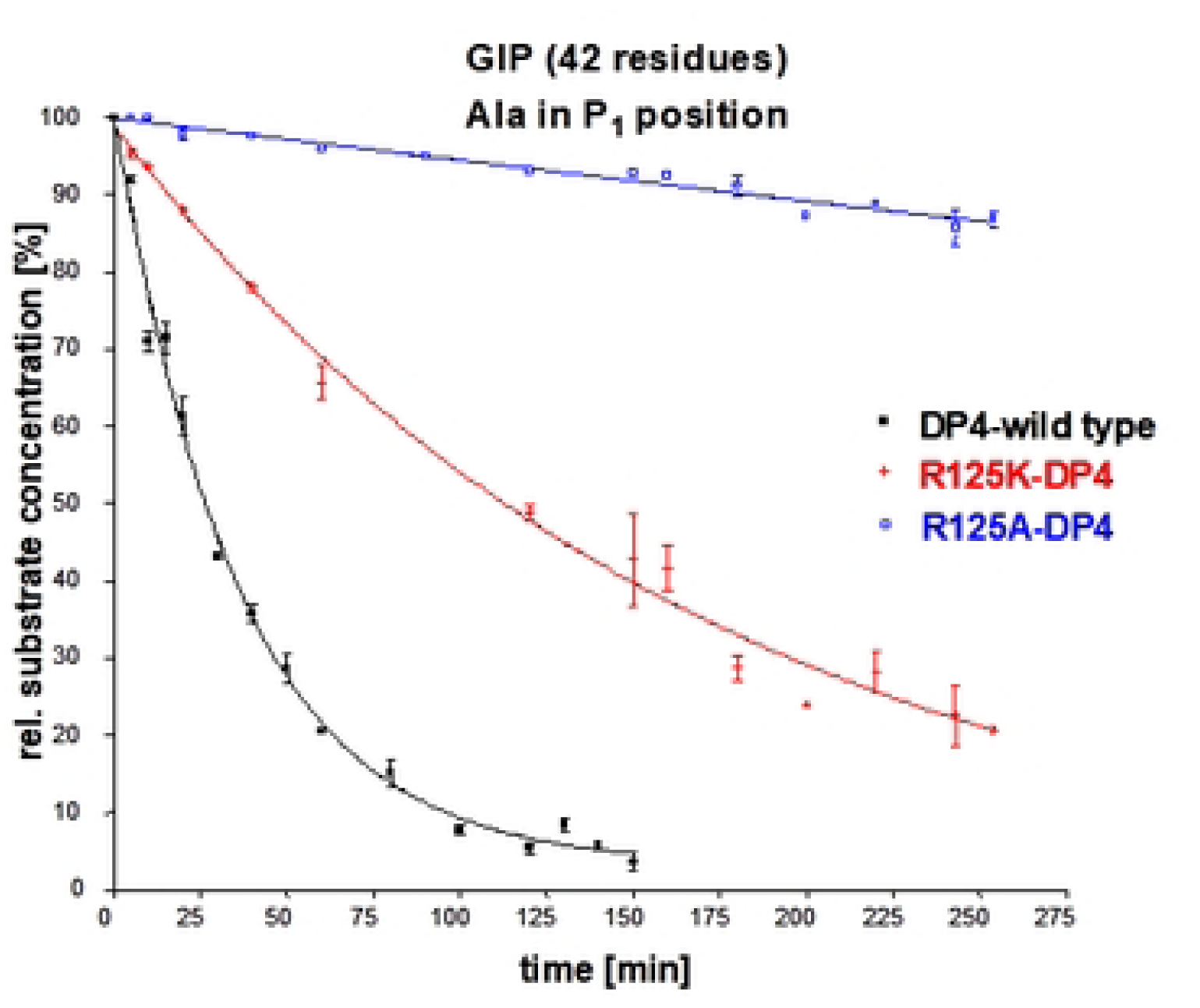

As shown in Table 5, the K_m_ as well as the k_cat_ values changed for the cleavage of GIP by R125K and R125A. Both enzyme variants displayed an impaired binding of NP Y while the k_cat_ values were not affected. The rate-limiting step for the hydrolysis of dipeptide derivative substrates with proline in P_1_ position is the deacylation reaction, for substrates with alanine in P_1_ position the acylation reaction [29]. Our results confirm that R125 interacts with the amino acid in P_1_’position. If the deacylation reaction is rate-limiting, R125 has no influence on this step, due to the fact that the C-terminal hydrolysis product (incl. amino acid in P_1_’position) is already released. In contrast, if the acylation reaction would be rate-limiting, R125 would contribute by the interaction with the carbonyl oxygen in P_1_’ to the coordination and the cleavage of the scissile bond. Since variation of R125 has no influence on the k_cat_ value for substrates with proline in P_1_ position (in our case NP Y and diprotin B) but for GIP (alanine in P_1_ position), it seems that the rate-limiting step for the hydrolysis of longer DP4 substrates is comparable with the kinetics of dipeptide derivatives, and depends whether proline or alanine is in P_1_ position [29].

Our investigations with the DP4 variants R125A and R125K further confirm the important role of the P_1_ amino acid in substrates. Processing of proline-substrates is still carried out after removing the side chain of R125, because the interaction between proline and the hydrophobic S_1_ pocket is still sufficient to enable substrate binding and catalysis. In contrast, processing of peptide hormones offering alanine (GIP) or serine (PACAP38) in P_1_ position is strongly affected if the side chain of R125 does not participate in their binding by replacement to alanine.

### Role of the glutamic acids E205 and E206 in substrate binding and catalysis

The enzymatic specificity of DP4 is usually dependent on the recognition of the P_1_ amino acid by the hydrophobic S_1_ binding pocket as well as the coordination of the positive charged N-terminus in P_2_ position. Crystal structures show, that binding of the substrate N-terminus occur by the negative charged glutamic acids 205 and 206 [31,32].

Our results confirm that the loss of the negative charge by variation of E205 and E206 significantly affects the substrate binding of dipeptide derivatives, tripeptides and peptide hormones. As previously described by Abbott and co-workers, the variant E206A has thereby stronger effects than E205A. Interestingly, both variants were able to cleave NP Y, and not GIP and PACAP38. This suggests also a significant role of the amino acid in P_1_ position. In case of NP Y, the recognition of proline by the hydrophobic S_1_ pocket is sufficient for proper orientation of the scissile bond to allow the nucleophile attack of the active serine. Thus, the coordination of the substrate N-terminus by both glutamic acids E205 and E206 is not essential for the hydrolysis of NP Y.

In summary, evaluated the importance of the amino acids R125, E205, E206 and N710 which are in close orientation to the catalytic triad in regard to their role in inhibition and binding of substrates with different length. It could be demonstrated that N710 is essential for processing substrates of all length while R125 supports the hydrolysis of peptide substrates consisting of more than two amino acids. Substrates displaying a proline in P_1_ position are well suited to fit perfectly into the hydrophobic S_1_ pocket by that strongly supporting their own catalysis. Alanine or serine in P_1_ position are no perfect matches for the S_1_ site. Nevertheless, many DP4 substrates offering smaller amino acids in P_1_ position, show comparable turnover rates to substrates with proline in P_1_. In that case, further stabilisation has to be provided by amino acids such as R125 and N710 or E205 and E206.

## Materials and Methods

### Strains and Plasmids

*P.pastoris* strain X-33 and the vector pPICZα C were purchased from Invitrogen (Thermo Fisher Scientific). *E.coli* XL-10 cells were provided by STRATAGENE EUROPE.

### Site directed mutagenesis

Based on the protein sequence of soluble hDP4, Δ1-36 hDP4, within the vector pPICZα C [36], site directed mutagenesis according to manual from Stratagene (QuickChange^TM^ Site-Directed Mutagenesis Kit) was carried out. In order to generate DP4 variants primers as listed in S1 Table were used.

### Transformation and expression of DP4

The *P. pastoris* expression constructs were transformed following the protocols from Invitrogen (Thermo Fisher Scientific).

A 2 l fermentation was inoculated with a pre-culture following the method described by Bär *et al.* [36]. The fermentation procedure was carried out in a 5 l Biostat-B reactor (B. Braun, Melsungen, Germany).

### Purification of DP4

Fermentation medium was centrifuged at 30,000 *g* for 20 minutes at 4°C to pellet the yeast cells. The supernatant was filtered to remove any residual solids and concentrated, using a tangential flow filtration system from Sartorius, Göttingen, Germany (cut-off: 30 kDA). Affinity chromatography was carried out at 4°C with a Ni-NTA sepharose column (Qiagen, Hilden, Germany). The column was pre-equilibrated with 50 mM NaH_2_PO_4_-buffer pH 7.6, 300 mM NaCl, 5 mM imidazole. The enzyme was eluted with increasing imidazole concentrations. The fractions with the highest DP4-content were pooled and concentrated to 0.5 ml using an Amicon ultrafiltration cell (cut-off: 10 kDA). This volume was applied to a superdex 200 gel filtration column (Cytiva life sciences). The column was equilibrated with 50 mM NaH_2_PO_4_-buffer pH 7.6 containing 300 mM NaCl with a flow rate of 0.25 ml/min. His(6)-37-766 hDP4 containing fractions were concentrated to 1 ml as described above.

### Determination of kinetic parameters

#### K_m_ determination of dipeptide derivatives and determination of IC_50_ values

The reactions were performed in 100 mM HEPES buffer, pH 7.6 at 30°C in a total assay volume of 270 µl by using 96-well plates. In order to measure K_m_ and k_cat_ values, the substrate concentrations varied from 4 x K_m_ to ¼ K_m_. The kinetic parameters were calculated with non-linear regression analysis. For IC_50_ determination the inhibitor concentrations varied from 10 x IC_50_ to 1/10 IC_50_, substrate Gly-Pro-AMC was used in the range of K_m_. Fluorescence was detected with a SPECTRAFluor Plus (TECAN) at an excitation wavelength of 380 nm and an emission wavelength of 465 nm. *K_m_ determination by LC/MS-MS assay.* Kinetic constants (*k_cat_* and *K_m_*) for the cleavage of the peptide hormones GIP and NP Y were determined by LC/MS-MS measurements. DP4 was incubated with different substrate concentrations in 10 mM Tris-buffer pH 7.6 at 30°C. Enzyme concentrations and incubation times were chosen to obtain a linear product formation. Reactions were stopped by addition of 19 volumes 0,1 % HCOOH. Cleavage products were quantified by LC/MS-MS assay.

#### Determination of cleavage products by MALDI-TOF mass spectrometry

Cleavage assays were performed in 10 mM Tris-buffer, pH 7.6. Enzyme concentration 5 x 10^-9^ M, substrate concentration 25 µM, reaction time 1-3 hours. Reactions were stopped by addition of 1 volume DHAP/DAHC-matrix, dissolved in 50 % (v/v) acetonitrile, after certain time points.

## Acknowledgements

We thank Nadine Jänckel and Anja Weber for technical assistance. We acknowledge support by the Federal Ministry of Education and Research (BMBF) under Grant No. 03FHP155AB.

## Supporting information

**S1 Fig. Chemical structure of Ile-thiazolidide**

**S1 Table. List of Primers.** All primers to perform side directed mutagenesis were purchased from metabion (Planegg/Steinkirchen, Germany).

## Notes

### Competing Interest Statement

The authors have declared no competing interest.

